# Pre-cuticle DPY-6 acts as a blueprint for aECM periodic organization in *C. elegans*

**DOI:** 10.64898/2026.02.21.707041

**Authors:** Sophie Mazzoli, Thomas Sonntag, Emma Cadena, Claire Valotteau, Susanna K. Birnbaum, Meera V. Sundaram, Nathalie Pujol

## Abstract

Apical extracellular matrices (aECMs) are essential for tissue integrity and function in multicellular organisms, but there is limited understanding of how such matrices are assembled and organized in the extracellular environment. The *Caenorhabditis elegans* cuticle, a model aECM that undergoes morphogenesis during each of the worm’s four larval molts, requires periodic circumferential furrows for structural integrity and immune regulation. Here, we show that furrow collagens must be cleaved from their N-terminal transmembrane domain for secretion and depend on the mucin-like pre-cuticle protein DPY-6 for their periodic assembly. While DPY-6 is dispensable for initial embryonic furrow formation, it acts as a mold during subsequent molts, ensuring pattern replication via its C-terminal cysteine cradle domain. These results reveal a central role for a transient matrix factor in organizing a complex periodically structured aECM.

**Author Summary:** In multicellular organisms, the extracellular matrix (ECM) provides structural support and regulates tissue function. Using the free-living worm Caenorhabditis elegans, we investigated how its apical ECM, the cuticle, forms a precise, repeating pattern of ridges called furrows. The cuticle is rebuilt at each of the worm’s four larval stages, providing a unique opportunity to study matrix morphogenesis in real time. We discovered that a transient protein, DPY-6, acts as a molecular mold to guide the self-organization of the matrix outside the epidermal cells. DPY-6 ensures that newly secreted proteins assemble into the correct periodic pattern during each rebuilding phase. Without DPY-6, the furrows lose their organization, leading to structural defects and immune system activation. Our findings reveal how a temporary scaffold can template the assembly of a complex, self-organizing structure. This work provides new insights into how biological matrices are built and maintained, with broader implications for understanding ECM assembly in health and disease.

## Introduction

All multicellular organisms rely on structured extracellular matrices to maintain tissue integrity and function. The *Caenorhabditis elegans* body cuticle is a rigid yet flexible apical extracellular matrix (aECM) structure that surrounds a single syncytial epidermal layer (reviewed in (Sundaram and Pujol, 2024)). It is a complex three-dimensional matrix composed of many different cuticle collagens and glycoproteins that are arranged in different layers and substructures. Therefore, the cuticle provides a powerful model for studying how extracellular matrices are assembled and organized.

One of the cuticle’s most striking features is the presence of circumferentially oriented furrows, which are distributed periodically along the length of the body (Adams et al., 2023; Cox et al., 1981; McMahon et al., 2003) (Fig. 1A). This periodic pattern is established during development, as embryos assemble the first larval cuticle (Costa et al., 1997), and a new cuticle is synthesized and shed at each larval stage through a process called molting (Lažetić and Fay, 2017). A transient pre-cuticle is assembled to help to pattern each new cuticle and to shed the old one (Sundaram and Pujol, 2024). The furrows of the new cuticle are precisely aligned to the furrow of the old cuticle (McMahon et al., 2003) (Fig 1A), and to some transient pre-cuticle components, like SPIA-1 and GRL-7 (Serra et al., 2024; Sonntag et al., 2025), but the mechanisms that first establish and then propagate this pattern are poorly understood.

**Fig 1.**
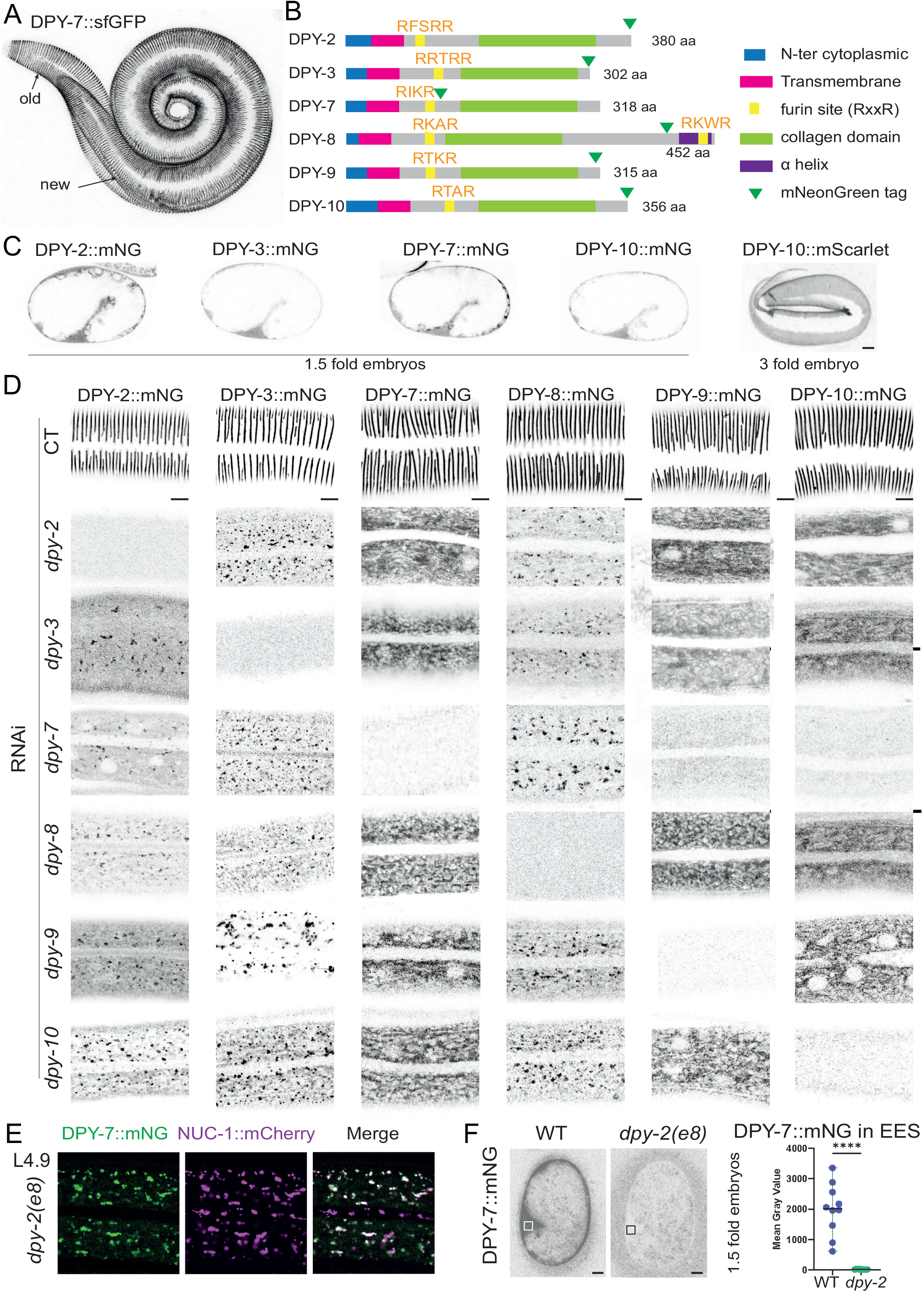
Furrow collagens are mutually dependent on each other for furrow assembly and secretion. (A) Maximum intensity projection from confocal image of DPY-7::sfGFP reporter in a molting L1 worm. Note the circumferential furrows in both the old and new cuticles. (B) Domain organization of the six furrow collagens proteins DPY-2, DPY-3, DPY-7, DPY-8, DPY-9 and DPY-10 in *C. elegans*, as annotated in InterPro (Mistry et al., 2021; Paysan-Lafosse et al., 2023) with the position of the mNeonGreen (mNG) tag insertion. (C) Representative confocal images of mNG tagged reporters for the 4 furrow collagens expressed in the 1.5-fold stage embryo DPY-2, DPY-3, DPY-7 and DPY-10. Images are shown in inverted grayscale and are representative of at least n = 10 worms per condition; scale bar, 5 µm. (D) Representative confocal images of mNG tagged reporters for all six furrow collagens in L4 worms during RNAi inactivation of our control *(sta-1)* and each *dpy* furrow collagen genes from L1 stage. In order to appreciate signal localization, focus is set on the cuticle layer for *sta-1* RNAi, whereas it is set in the epidermis for *dpy* RNAi, as no signal was never observed at the cuticle level. Note that the expression levels are not comparable between strains and different sub-stages are presented from early to mid L4 stage. Images are shown in inverted grayscale and are representative of at least n = 10 worms per condition; scale bar, 5 µm. (E) Representative confocal images of DPY-7::mNG (green) and NUC-1::mCherry (magenta) in L4.7 worm and the overlay of both signals ; n>10; scale bar, 5 µm. (F) Reduced DPY-7 secretion in *dpy-2(e8)* mutant embryos. Images are shown in inverted grayscale; scale bar, 5 µm. The graph shows mean fluorescence intensity in the extra-embryonic space (EES) of 1.5-fold stage embryos, corresponding to the boxed regions in the images at left; n=10 per each genotype.

Mutants lacking periodic furrows have proven invaluable for studying the interplay between cuticle structure and epidermal homeostasis. Mutations in any of the six furrow collagens—DPY-2, DPY-3, DPY-7, DPY-8, DPY-9, and DPY-10—result in the absence of periodic furrows (Cox et al., 1980; McMahon et al., 2003; Thein et al., 2003). These mutations also lead to persistent immune activation (PIA), a response akin to that triggered by molting, physical injury, or infection (Aggad et al., 2023; Dodd et al., 2018; Pujol et al., 2008a; Pujol et al., 2008b; Taffoni et al., 2020; Zugasti et al., 2014). This dual phenotype— loss of structural organization and immune dysregulation—suggests that furrow collagens play a critical role in both structuring the cuticle and modulating defense responses in the epidermis.

Given the importance of furrow collagen organization for cuticle integrity and epidermal communication, we sought to investigate where and when these collagens are expressed, how they are secreted into the cuticle, and what molecular scaffolds guide their assembly. One candidate scaffold is DPY-6, a mucin-like protein that was previously proposed to be involved in cuticle deposition (Sun et al., 2022). It contains a conserved Cysteine cradle domain (CCD) shared with five other *C. elegans* proteins predicted to be secreted into the cuticle (Sonntag et al., 2025). The structured core of this domain, stabilized by conserved cysteine interactions, suggests it may mediate homomeric and heteromeric interactions (Sonntag et al., 2025). Notably, DPY-6 is one of the first secreted proteins marking the initiation of each new cycle of aECM morphogenesis (Meeuse et al., 2020; Sonntag et al., 2025), positioning it as an ideal candidate for scaffolding the new matrix.

In this study, we demonstrate that furrow collagens must be cleaved from their N-terminal transmembrane domain to be secreted from the syncytial epidermis into the cuticle. Once secreted, they require DPY-6 to assemble into a periodic circumferential pattern. We show that DPY-6 is a transient (pre-cuticle) matrix component not present in mature furrows. While the first cuticle built in the embryo does not require DPY-6 for this periodic alignment, we show that DPY-6 is essential as a mold to replicate the position of the periodic furrow at each molt. The recruitment of new collagens to the furrow depends on the CCD of DPY-6. These findings suggest that the DPY-6 mucin-like protein scaffolds the assembly of a periodically organized extracellular matrix—a structure essential for both physical protection and immune homeostasis.

## Results

### Furrow collagens assemble into periodic structures after secretion

Previous studies have shown that DPY-2, DPY-3, DPY-7, DPY-8, DPY-9, and DPY-10 are cuticular collagens specifically localized to the furrows (reviewed in (Sundaram and Pujol, 2024)). We generated two new knock-in (KI) strains for DPY-7 and DPY-8 and collected existing strains for the four others (Ragle et al., 2026) (Fig 1B). We confirmed that all six collagens localize to the furrows, either from the 3-fold embryo stage for DPY-2, DPY-3, DPY-7, and DPY-10, or from mid-L1 for DPY-8 and DPY-9 (Fig 1C&D & S1A). Previous antibody staining studies showed that DPY-7 is detected intracellularly from the comma stage but only associates with the cuticle at the 3-fold embryo stage (McMahon et al., 2003). We confirmed and extended these findings to all early furrow collagens. In young (1.5-fold) embryos, when the first pre-cuticle is present, these collagens are initially broadly secreted and found between the eggshell and the embryo, before gradually becoming patterned into furrows during later stages of cuticle assembly (Fig 1C). These observations are similar to those reported for other non-furrow collagens (Adams et al., 2023; Birnbaum et al., 2023), consistent with the model that patterning occurs at a post-secretory step.

### Furrow collagens are mutually dependent on each other for secretion

It has been previously suggested that DPY-2, DPY-3, DPY-7, DPY-8, DPY-9, and DPY-10 function together. The removal of any one of these six collagens results in the loss of periodic furrows and a dumpy (Dpy) body shape, but also in other phenotypes in the underlying epidermis that are characteristic of furrow loss and not found in other Dpy mutants, like persistent immune activation (Pia), osmotic stress resistance due to glycerol accumulation (Osm), and loss of transient cytoskeleton alignment (Aggad et al., 2023; Dodd et al., 2018; McMahon et al., 2003; Sonntag et al., 2025; Wheeler and Thomas, 2006). Additionally, these collagens have been suggested to be required for each other’s secretion (McMahon et al., 2003; Sonntag et al., 2025), similar to what has been described for another pair of collagens, SQT-3 and DPY-17 (Birnbaum et al., 2023; Novelli et al., 2006). We tested the inactivation of each of the furrow collagen genes on the secretion of all six mNG-tagged furrow collagens (Fig 1D). In all cases, none of the furrow collagens were secreted and present at the cuticle level. Instead, they were retained in the cytoplasm, either in a halo pattern around the epidermal nuclei, suggesting retention in the endoplasmic reticulum, or present in vesicles within the epidermis. This pattern was also observed in a *dpy-2(e8);* DPY-7::mNG FBN-1::mCherry mutant strain, where DPY-7::mNG was found in similar vesicles as the pre-cuticular component FBN-1 in late L4 (Fig S1B) (Balasubramaniam et al., 2023; Kelley et al., 2015). We confirmed that DPY-7::mNG was found in similar vesicles as the lysosomal hydrolase NUC 1::mCherry that served as an endosomal and a lysosomal reporter, as previously described (Li et al., 2016; Miao et al., 2020) (Fig 1E) suggesting that in a *dpy-2* mutant, DPY-7::mNG is targeted to lysosome for degradation instead of being secreted. Moreover, in *dpy-2(e8)* mutant 1.5 fold embryos, DPY-7::mNG was no longer secreted into the space between the eggshell and the embryo (Fig 1F). These data support the model that DPY-2, DPY-3, DPY-7, DPY-8, DPY-9, and DPY-10 travel together through the secretory pathway.

### Secretion of furrow collagen requires cleavage from their N-ter transmembrane domain

Approximately two thirds of *C. elegans* cuticle collagens are predicted to be secreted in type II orientation, featuring a cytosolic N-terminus followed by a transmembrane (TM) domain (Sundaram and Pujol, 2024; Teuscher et al., 2019). Similar to mammalian MACITs (membrane-associated collagens with interrupted triple helices) and other collagen-related transmembrane proteins (Tu et al., 2015; Wakabayashi, 2020), these transmembrane cuticle collagens may remain associated with cell surfaces or be released into the matrix through proteolysis. All six furrow collagens possess a TM domain followed by a potential furin cleavage site (Fig 1B).

To test whether furrow collagens are correctly secreted and positioned in the matrix while retaining their attachment to the epidermis through their TM domain, we N-terminally tagged DPY-7 and DPY-8 (mNG::DPY-7 & mNG::DPY-8) before the TM domain and compared them with the C-terminally tagged versions (DPY-7::mNG and DPY-8::mNG) (Fig 2A). We never observed secretion of the N-terminally tagged versions of DPY-7 or DPY-8. Instead, we observed accumulation in vesicles within the epidermis, with larger and denser vesicles at the mid-L4 stage, when furrow collagens expression are known to be at their peak in oscillation (Meeuse et al., 2020; Sonntag et al., 2025) (Fig 2C&D). These vesicles are similar to the NUC 1::mCherry lysosomal vesicles (Fig 2E). Since these animals exhibited, however, a wild-type length and not a Dpy phenotype (Fig 2B), this suggests that the N-terminally tagged proteins are functional and potentially cleaved after the TM domain to reach their correct position in the cuticle.

**Fig 2.**
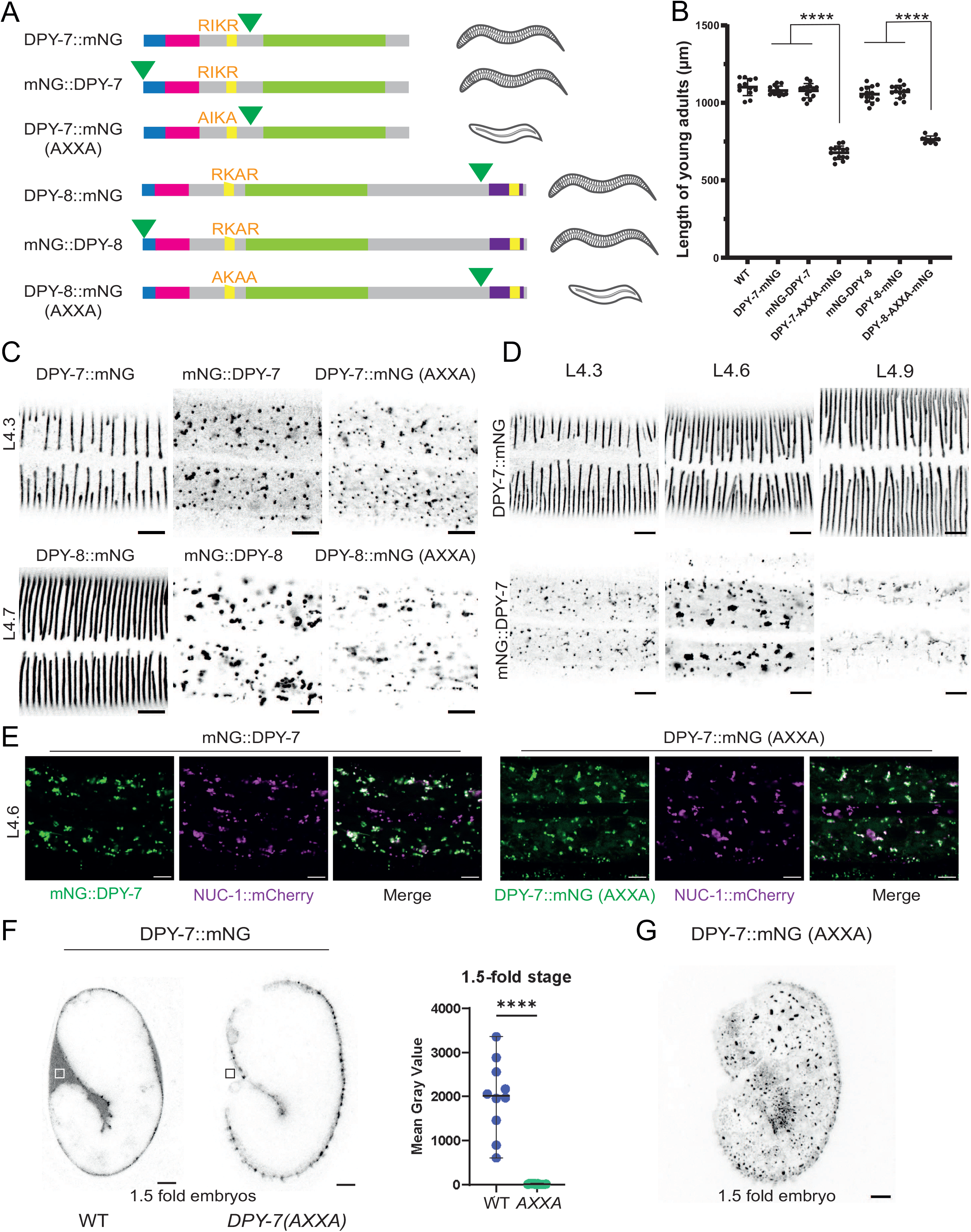
Secretion of furrow collagens requires cleavage from their N-ter transmembrane domain. (A) Schematic representations of positions of the insertion of the mNG tag for DPY-7 & DPY-8 strains, with sequence of each furin site, either wild-type RXXR or mutated AXXA, removing both Arginine known to be required for furin-dependent cleavage (Seidah and Prat, 2012). Schematic representation of the observed phenotypes (wild-type or dumpy) for each strain (B) Worm lengths measured at young adult stage; n>10; \*\*\*\**p* < 0.0001. (C) Representative confocal images of mNG signal pattern in L4 larvae. In order to appreciate signal localization, the focus is set on the cuticle layer for DPY-7::mNG and DPY-8::mNG strains, whereas it is set in the epidermis for the others, as the collagen are not secreted into the cuticle. Images are shown in inverted grayscale; n>25 for DPY-7 tagged strains and n>5 for DPY-8 tagged strains; scale bar, 5 µm. (D) Representative confocal images of a timeline during the L4 stage for DPY-7::mNG and mNG::DPY-7 strains. In order to appreciate signal localization, focus is set on the cuticle layer for DPY-7::mNG, whereas it is set in the epidermis for mNG::DPY-7. Images are shown in inverted grayscale; n>5 worms per stage; scale bar, 5 µm. (E) Representative confocal images of mNG (green), NUC-1::mCherry (magenta) and merged signals in the epidermis of L4 larvae showing that both the N-cleaved mNG::DPY-7 and the DPY-7::mNG(AXXA) are in similar vesicle as NUC-1::mCherry; n>25; scale bar, 5 µm. (F) Retention of DPY-7 in *DPY-7(AXXA)* mutant embryos. Images are shown in inverted grayscale. Scale bar, 5 µm. The graph shows mean fluorescence intensity in the extra-embryonic space (EES) of 1.5-fold stage embryos, corresponding to the boxed regions in the images at left; n=10 per each genotype. (G) Representative maximum intensity projection from confocal image of DPY-7::mNG reporter in 1.5-fold stage *DPY-7(AXXA)* mutant embryo; scale bar, 5 µm.

To test whether the furrow collagens are cleaved at their potential furin site after the TM domain, we used CRISPR/Cas9 to mutate the predicted RXXR motif to AXXA within the endogenous DPY-7::mNG and DPY-8::mNG fusions. Similar mutations have previously been shown to abrogate furin cleavage in other collagens, such as SQT-3 and DPY-17 (Birnbaum et al., 2023). In the DPY-7::mNG(AXXA) and DPY-8::mNG(AXXA) mutants, as observed with the N-terminally tagged proteins, the collagens were never secreted into the matrix but were retained in vesicles within the epidermis (Fig 2C). These vesicles are similar to the NUC 1::mCherry lysosomal vesicles (Fig 2E) suggesting that the furin cleavage mutant protein DPY-7::mNG(AXXA) is targeted to lysosome for degradation instead of been secreted. This absence of secretion and retention was also observed in the early embryo (Fig 2F-G). Moreover, the worms exhibited a Dpy phenotype (Fig 2B), suggesting that the non-cleaved proteins cannot fulfil their function in the matrix. Together, these results are consistent with a model in which furrow collagens are cleaved to allow their correct secretion.

### The proprotein convertase BLI-4 is partially required for furrow collagen secretion

The convertase BLI-4 has previously been implicated in the N-terminal processing of multiple cuticle collagens, including SQT-3 and DPY-17 (Birnbaum et al., 2023; Thacker et al., 1995). Since *bli-4* null mutants are lethal and full RNA inactivation also results in L1 lethality (Fig S2A), directly testing whether BLI-4 is required for specific collagen cleavage is challenging. To circumvent this, we used RNA inactivation at the L3 stage and examined at the L4 stage the newly formed adult cuticle, which develops underneath the L4 cuticle (Fig 3A). Partial *bli-4* inactivation resulted in misalignment of the furrow collagen DPY-8::mNG in the newly formed adult cuticle, while collagens remained aligned in the L4 cuticle formed before *bli-4* inactivation (Fig 3B-D). Additionally, we observed accumulation of non-secreted collagen in vesicles within the epidermis (Fig 3C&D). When we applied a 3-hour pulse of *bli-4* inactivation at the L4 stage and examined the next generation, approximately one-fifth of the population exhibited a severe Dpy phenotype, yet these worms reached adulthood and produced wild-type progeny (Fig S2B). These Dpy worms displayed disorganized furrow collagen in the cuticle. Interestingly, in regions between abnormally branched seam cells with duplicated alae, the collagen aligned in a more longitudinal orientation, as if constrained by the abnormal seam boundaries (Fig 3E; Fig S2C). To further investigate the requirement of BLI-4, we used previously described hypomorphic and knockout mutations (Birnbaum et al., 2023). In *bli-4(cs302)* hypomorphic larvae, we observed the same phenotype of disorganized furrow collagens using the DPY-7::sfGFP strain (Fig 3F). Additionally, when examining assembly of the first cuticle in embryos, initial DPY-7::sfGFP secretion was severely reduced in *bli-4(cs281)* null mutants compared to WT (Fig 3G), although over time some DPY-7::SfGFP appeared to join the cuticle in a disorganized furrow-like fashion (Fig 3H). The other furin protease known to be expressed in the epidermis is KPC-1. Inactivation of *kpc-1* alone didn’t give any phenotype and double RNAi of kpc-1 and bli-4 did not aggravate the phenotype of *bli-4* (Fig S2D).

**Fig 3.**
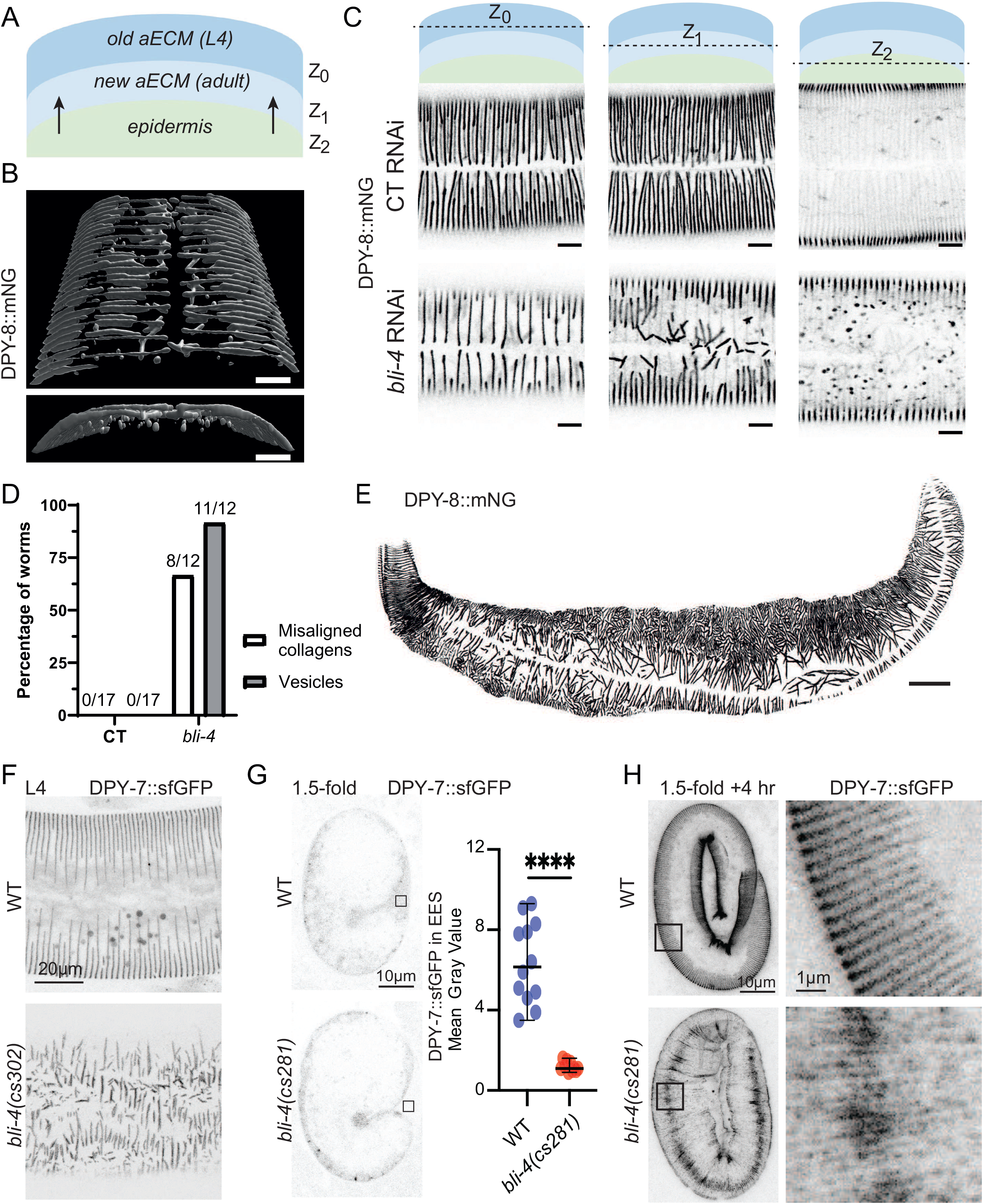
The proprotein convertase BLI-4 is partially required for furrow collagen secretion. (A) Schematic representation of the worm morphology at the cuticle level. The old cuticle (dark blue) will be replaced by the new one (light blue), that is built by components coming from the epidermis (green). (B-D) L3 larvae of DPY-8::mNG were put on *sta-1* or *bli-4* RNAi until L4 stage; (B) 3-dimensional reconstitution of a *bli-4* RNAi L4 larvae (same worm as C); scale bar, 30 µm. (C) Representative confocal images of DPY-8::mNG reporter in late L4 larvae (same worm as B), at different levels (old cuticle, new cuticle and epidermis); n>10. Images are shown in inverted grayscale; scale bar, 5 µm. (D) Quantification of observed DPY-8::mNG patterns. Misaligned collagens are observed at the Z_1_ level (new cuticle) and vesicles at the Z_2_ level (epidermis). (E) Dpy L4 larva progeny from a DPY-8::mNG mother subjected to a 3 hour pulse of *bli-4* RNAi during the L4 stage (see also Fig. S2). Representative maximum intensity projection from confocal images in inverted grayscale; n>5; scale bar, 30 µm. (F-H) Representative confocal images of DPY-7::SfGFP *(qxIs722)* in *bli-4* mutants (n=10 or more each). (F) Disorganized furrows in *bli-4(cs302*) hypomorphs that survive to a late larval stage (48 hours after egg lay). (G) Reduced DPY-7 secretion in *bli-4(cs281*) null mutant embryos. The graph shows mean fluorescence intensity in the extra-embryonic space (EES) of 1.5-fold stage embryos, corresponding to the boxed regions in the images at right. (H) Confocal projections of older embryos, with boxed regions enlarged at right. Some DPY-7 furrow-like signal is detected in the cuticle of *bli-4(cs281*) mutants, suggesting residual secretion, possibly due to maternal rescue or another compensating protease.

While mutation of the furin cleavage site in DPY-7::mNG(AXXA) and DPY-8::mNG(AXXA) completely blocks furrow collagen secretion, these mutants remain fully viable. In contrast, *bli-4(0)* null mutants or RNAi-treated worms exhibit embryonic lethality, likely due to cumulative effects on multiple substrates, including essential embryonic collagens such as SQT-3 and DPY-17 (Birnbaum et al., 2023). Despite its broader role in processing multiple substrates, our findings suggest that BLI-4 is specifically required for the cleavage of furrow collagens and may act redundantly with another furin-like protease to ensure their efficient secretion.

### DPY-6 is a mucin-like CCD protein localized early and transiently at the periodic furrows

The *C. elegans* epidermis is a giant syncytium without junctional landmarks, apart from the longitudinal seam cell, raising the question: what directs the circumferential periodic organization of furrow collagens once secreted into the cuticle? The mucin-like protein DPY-6, containing a conserved Cysteine cradle domain (CCD) (Sonntag et al., 2025) emerged as a compelling candidate due to several key features. First, *dpy-6* gene expression is oscillating and its peak precedes that of known pre-cuticular and furrow collagen genes (Fig. 4A) (Meeuse et al., 2020); indeed, we previously suggested that DPY-6 marks the onset of a new cycle at the end of each molt (Sonntag et al., 2025). Second, DPY-6 localizes to the furrows (Sonntag et al., 2025), as confirmed in a strain co-expressing DPY-6::mNG and DPY-10::mScarlet, with DPY-6 found in between the new and the old L4 cuticle in late L4 stage (Fig. 4B).

**Fig 4.**
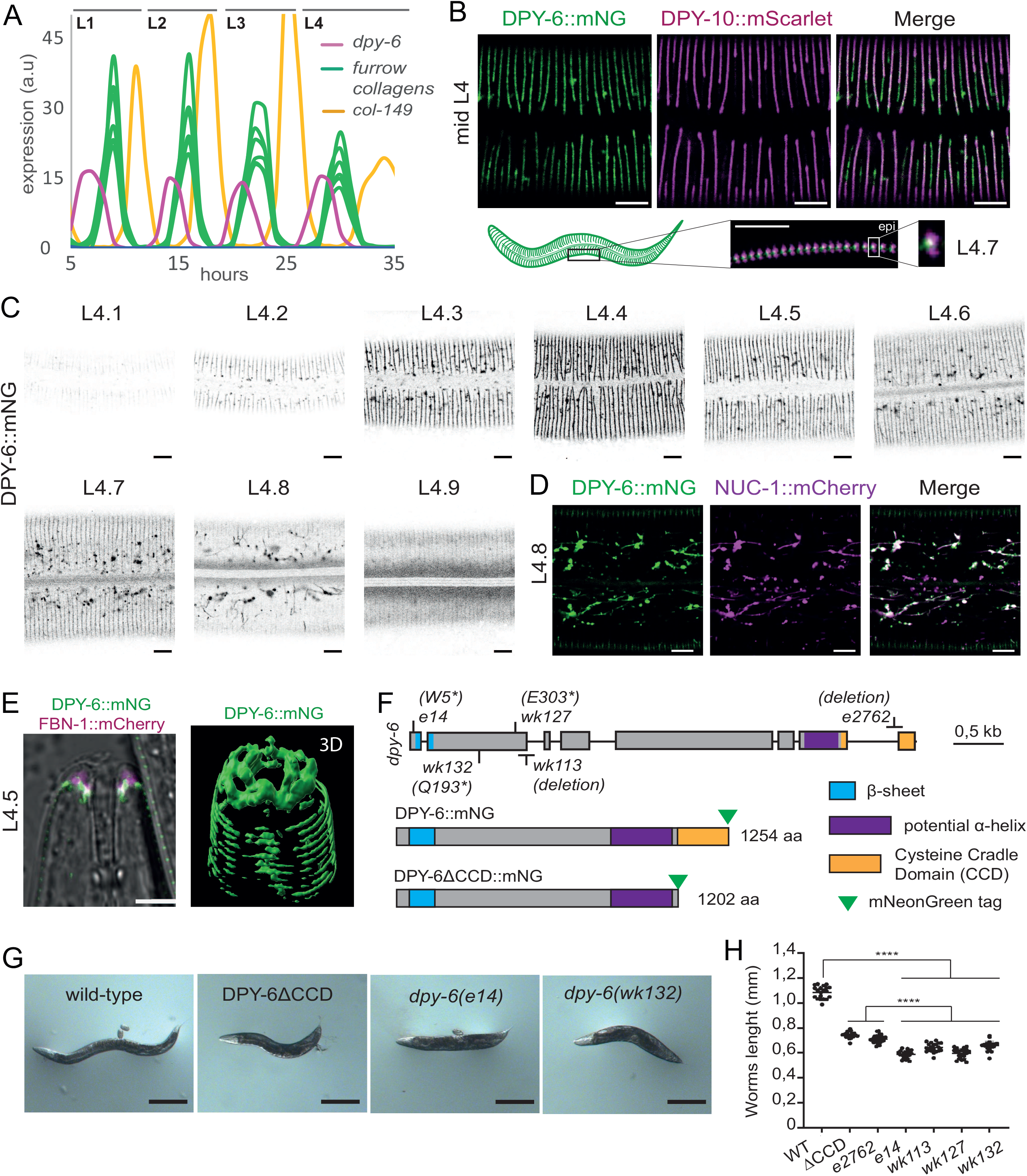
DPY-6 localizes to furrows and its gene expression is oscillatory between each molt. (A) *dpy-6* gene expression oscillates between each molt, peaking before the genes encoding furrow collagens and a late expressed collagen *col-149*. Data reanalyzed from (Meeuse et al., 2020). (B) Representative confocal images of DPY-6::mNG (green) and DPY-10::mScarlet (magenta) in a mid-L4 worm and the overlay of both signal; n=15; scale bar, 5 µm. Zoom of L4.7 larvae’s side showing DPY-10::mScarlet in both old and new cuticles, and DPY-6::mNG in between. scale bar, 5 µm (C) Representative confocal images of DPY-6::mNG during L4 stage, from L4.1 to L4.9, using vulval shape as a proxy for developmental timing (Mok et al., 2015), shown in inverted grayscale; n>5 for each sub-stage; scale bar, 5 µm. (D) Representative confocal images of DPY-6::mNG (green) and NUC-1::mCherry (magenta) in L4.8 worm and the overlay of both signals ; n=30; scale bar, 5 µm. (E) Left, representative confocal images of the overlay of DPY-6::mNG (green), FBN-1::mCherry (magenta) and the transmission channel in mid-L4 worm mouth; n=5; scale bar, 5 µm. Right, 3D reconstruction of the DPY-6::mNG signal. (F) Schematic representation of the *dpy-6* gene, with its associated mutant alleles. In addition to the previously characterized *e14* allele (W5*), four additional alleles have been sequenced: *wk127 (E303*), wk132 (Q193*), wk113* (hundred base pairs deletion in the second exon leading to a premature STOP) and *e2762* (61 base pairs deletion in the 7^th^ intron and the last exon, leading to mis-splicing and so, unfunctional CCD). Schematic representation of DPY-6 and DPY-6ΔCCD mutated proteins with domain organization as annotated in InterPro and the mNG tag at the C-ter. (G) Representative images of wild-type, *dpy-6(e14), dpy-6(wk127)* and DPY-6ΔCCD mutant young adult worms; n>50; scale bar, 100 µm. (H) Worm lengths measured at young adult stage between different *dpy-6* mutant alleles and the wild-type; n=20; ****p < 0.0001

We further demonstrated that DPY-6 is a bona fide pre-cuticular component, transiently expressed in the apical matrix. Using vulval shape as a developmental timing proxy (Mok et al., 2015; Sonntag et al., 2025), we observed that DPY-6::mNG begins to mark the furrows as early as the L4.2 stage, peaks at L4.4, and is subsequently removed from the furrows, accumulating in vesicles similar to NUC-1::mCherry lysosomes at later stages (Fig. 4C-D). This trafficking pattern aligns with previously described pre-cuticular components (Forman-Rubinsky et al., 2017; Katz et al., 2022; Sonntag et al., 2025; Sundaram and Pujol, 2024).

DPY-6::mNG was also found transiently lining the beginning of the mouth during the mid-L4 stage (Fig 4E). DPY-6::mNG signal was found mainly colocalizing and just below FBN-1::mCherry signal in L4.5 stage. FBN-1 is a ZP-fibrillin known to be required for the adhesion of the pharynx to the cuticle (Balasubramaniam et al., 2023; Kelley et al., 2015). This is in accordance with the previously characterized role of DPY-6 in mouth formation (Sun et al., 2022). Collectively, these results demonstrate that DPY-6 is a typical pre-cuticle component, as its signal peaks during the intermolt period at the onset of new cuticle synthesis, after which it is cleared by endocytosis.

### Removal of the cysteine cradle domain is sufficient to give a DPY phenotype

The *dpy-6* gene was first identified based on its Dpy (Dumpy) phenotype, characterized by shorter and wider adult worms compared to wild-type animals, similar to the phenotype observed in furrow collagen mutants (Brenner, 1974). Subsequent studies isolated additional *dpy-6* alleles including *wk113, wk127* and *wk132*, as suppressors of the elongated body size phenotype in *lon-2* mutants (Lakdawala et al., 2019). Sequencing of these alleles revealed premature stop codons in the early coding region of the gene, potentially generating null alleles (Fig. 4E).

DPY-6 is a largely disordered protein, featuring a conserved Cysteine Cradle Domain (CCD) at its C-terminus, which is predicted to form a hydrophobic pocket in the shape of a cradle, potentially binding a partner(Sonntag et al., 2025). Structural predictions also indicate a coil coiled domain adjacent to the CCD and a beta-sheet within the N-terminal domain (Fig. 4F). To investigate the functional importance of the CCD, we generated a deletion in the DPY-6::mNG-tagged strain, referred to as DPY-6ΔCCD::mNG (Fig. 4F). DPY-6ΔCCD::mNG mutant worms exhibited a shorter body length compared to wild-type animals, though not as severe as the four potential null alleles (Fig. 4F-H). Furthermore, we found that the previously reported *dpy-6(e2762)* hypomorphic mutant (Davis et al., 2022) harbors a deletion in the CCD (Fig 4F) and has a size similar to DPY-6ΔCCD::mNG mutants (Fig 4H). This suggests that while the CCD is essential for full DPY-6 function, the remaining protein retains partial activity.

### DPY-6 acts as a blueprint for furrow collagens

Topological imaging by atomic force microscopy (AFM) revealed the loss of the periodic organization of the furrows in *dpy-6(e14)* young adult worms, with adjacent furrows appearing to point toward each other (Fig. 5A). Upon closer examination of the furrow collagens, we found that genetic ablation or RNAi-mediated knockdown of DPY-6 results in a loss of periodic organization of all six furrow collagens; the collagens are still secreted in the cuticle but are found associated in mis-oriented large bundles (Fig 5B&C). Notably, deletion of the C-terminal cysteine cradle domain (CCD) alone is sufficient to recapitulate this phenotype (Fig. 5D). Interestingly, DPY-6ΔCCD::mNG is still secreted in the cuticle albeit at low levels and colocalizes with the disorganized furrow collagen DPY-10::mScarlet (Fig 5D). These data indicate that DPY-6 patterns cuticle furrows and requires its CCD domain to do so. Furthermore, DPY-6 can associate with furrow collagens via a domain other than the CCD.

**Fig 5.**
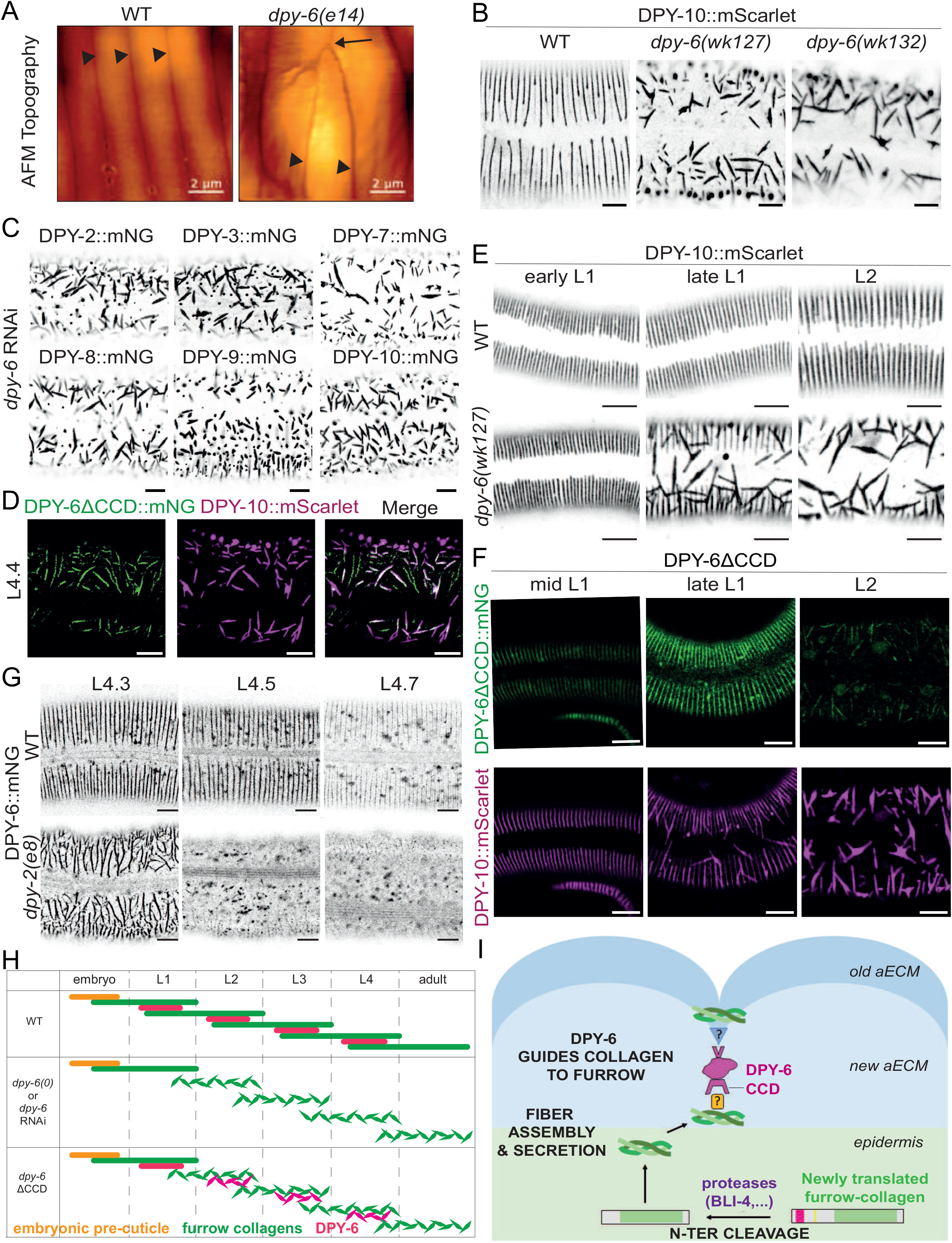
DPY-6 acts as a blueprint for furrow collagens. (A) AFM topography of the cuticle in wild-type or *dpy-6(e14)* young adult worms. Arrowhead are pointing to furrows and the arrow is pointing to their intersection; n>5; scale bar, 2 µm. (B) Representative confocal images of DPY-10::mScarlet in wild-type, *wk127* and *wk132* mid-L4 worms. Image are shown in inverted grayscale; n=15; scale bar, 5 µm. (C) RNAi inactivation of *dpy-6* gene from L1 stage in the six furrow collagen mNG-tagged strains prevents furrow-collagen alignment in the L4 stage; Representative confocal images, shown in inverted grayscale; n>10; scale bar, 5 µm. (D) Representative confocal images of DPY-6ΔCCD::mNG (green), DPY-10::mScarlet (magenta) and the overlay of both signals in mid-L4 worm; n=15; scale bar, 5 µm. (E) Representative confocal images of DPY-10::mScarlet in wild-type or in *wk127*, from early L1 to L2 staged worms. Stages have been defined using gonad development. Image are shown in inverted grayscale; n>10; scale bar, 5 µm. (F) Representative confocal images of DPY-10::mScarlet (magenta) in DPY-6ΔCCD::mNG (green) worms, stages going from early L1 to L2. Stages have been defined using gonad development; n>10; scale bar, 5 µm. (G) Representative confocal images of DPY-6::mNG (green) in L4 larvae at different sub stages, either in wild-type or in *dpy-2(e8)* mutant; Images are shown in inverted grayscale; n>10; scale bar, 5 µm. (H) Schematic time line of the furrow collagens in green and DPY-6 in magenta during the four larval stages in wild-type and in different *dpy-6* inactivation conditions. Orange bar indicates unknown factor(s) that pattern the furrows in the first cuticle. (I) Cartoon presenting the proposed model for furrow collagen organization. Furrow collagens are cleaved and then secreted together in the aECM. DPY-6 localizes to the existing furrows and acts as blueprint for positioning the new collagens. The Cysteine-Cradle Domain of DPY-6 interacts (directly or indirectly) with furrow collagens of the newly forming cuticle. We propose that DPY-6 uses a different domain (upper V-shape) to interact (directly or indirectly) with the furrow collagens present in mature furrows, thereby positioning new furrow collagens directly below the old ones.

To determine whether DPY-6 associates with furrow collagens from the old or new cuticle, we examined the first L1 cuticle, which is synthesized during embryonic development (Sundaram and Pujol, 2024). Four of the six furrow collagens—DPY-2, DPY-3, DPY-7, and DPY-10— are expressed embryonically and assemble into L1 cuticle furrows (Fig. 1 & Fig S1). DPY-6 is not visible in the embryo nor in newly hatched L1s, but it starts to be observed in the mid L1 stage, as the L2 cuticle is being built (Fig S1A). Moreover, in a *dpy-6(wk127)* null mutant, the DPY-10::mScarlet pattern in newly hatching L1s was comparable to the wild-type but started to be non-periodic as soon as the L2 cuticle began to form in the mid L1 (Fig 5E). This suggests that furrow collagens can assemble embryonically without DPY-6 but require DPY-6 to replicate their periodic pattern in larvae.

Interestingly, in the DPY-6ΔCCD::mNG mutant strain, DPY-6ΔCCD::mNG initially localizes to the furrow in a periodic pattern in L1 larvae, but the furrow collagens of the L2 cuticle fail to align with this pattern (Fig. 5F), similar to what is seen in *dpy-6* null mutants. This suggests that DPY-6 can associate with furrows of the old (L1) cuticle via one of its other domains, but the CCD is essential for positioning newly secreted furrow collagens into the periodic pattern (Fig 5H&I). Interestingly, in a *dpy-2(e8)* mutant, which lacks normal parallel furrows, DPY-6::mNG is disorganized in randomly oriented stripes in the cuticle at the beginning of the L4 stage, and then rapidly removed for degradation in the epidermis (Fig 5G). Together, these observations indicate that DPY-6 acts as a molecular mold, propagating the periodic patterning of furrow collagens to the next cuticle.

## Discussion

The *C. elegans* cuticle is a highly organized apical extracellular matrix (aECM) that undergoes periodic remodeling during each larval molt. In this study, we reveal that the pre-cuticle protein DPY-6 acts as a molecular scaffold, orchestrating the periodic assembly of furrow collagens and ensuring the propagation of this pattern across successive molts. Our findings provide mechanistic insights into how a transient, pre-cuticular component can template the organization of a complex ECM, with broader implications for understanding ECM assembly and function in multicellular organisms.

### Furrow Collagens: Cleavage, Secretion, and Assembly

Collagens in *C. elegans* undergo a regulated process of cleavage, secretion, and extracellular assembly, essential for their incorporation into the cuticle structure. Furrow collagens are predicted to be synthesized as type II transmembrane proteins, featuring a short cytosolic N-terminal domain, a transmembrane (TM) helix, and an extracellular collagenous C-terminus. This transmembrane orientation is reminiscent of mammalian MACITs, such as collagen XIII, which also require proteolytic cleavage for proper function (Tu et al., 2015; Wakabayashi, 2020). The presence of a conserved furin cleavage site (RxxR) immediately following the TM domain in all six furrow collagens suggests that they are processed by proprotein convertases, such as BLI-4, a *C. elegans* furin-like enzyme known to cleave multiple cuticle collagens (Birnbaum et al., 2023; Thacker et al., 1995). This cleavage is likely a rate-limiting step for secretion, as mutations in the furin site result in the absence of secretion and block the incorporation into the cuticle as was found for other embryonic collagens (Birnbaum et al., 2023). This process is reminiscent of other ECM proteins, such as mammalian collagens, which require proteolytic processing before forming higher-order structures (Revell et al., 2021).

The mutual dependence of furrow collagens for secretion suggests that these proteins associate intracellularly, potentially forming heteromeric complexes prior to secretion. Collagens are known to form heterotrimers, and these collagen triple helices are known to further assemble into bundles reminiscent of alpha-helical coiled-coils; the bundling of collagen triple helices remains, however, poorly understood (Revell et al., 2021). A recent study demonstrated that six triple helices of collagen can assemble into a discrete supramolecular structure (Yu et al., 2023), suggesting that a similar higher-order multimeric arrangement could underlie the interdependence observed among the six furrow collagens. This interdependence reflects a requirement for coordinated trafficking or stabilization within the secretory pathway. Whether these collagens form a supramolecular structure during secretion remains an open question for future investigation. The retention of unsecreted collagens in vesicles highlights the quality control mechanisms that degrade misfolded or improperly processed ECM components (Miao et al., 2020). This suggests that the assembly of furrow collagens into a secretion-competent complex is tightly regulated, and any disruption leads to their degradation. Future structural and biochemical studies will be needed to determine the exact stoichiometry and interactions among these collagens, as well as whether additional factors (e.g., chaperones or assembly proteins) are required for their proper folding and secretion.

### DPY-6 as a Transient Molecular Mold for Periodic ECM Organization

Once secreted into the extracellular space, furrow collagens must assemble into higher-order structures at the proper location. Our findings establish that DPY-6 serves as a molecular template to replicate the periodic pattern of furrow collagens (Fig. 5I). While DPY-6 is not required for collagen secretion or fibril formation per se—as furrow collagens still assemble into fibrils or bundles in its absence—these structures are no longer periodically organized. The Cystein Cradle Domain is critical for recruiting newly secreted collagens to the furrow, as its deletion disrupts fiber alignment in the L2 cuticle. However, DPY-6ΔCCD::mNG still localizes to furrows in L1 larvae, suggesting that another domain— potentially in the N-terminal region (Li et al., 2026)—anchors DPY-6 to the old furrows. By engaging distinct domains to interact with existing furrows and newly secreted collagens, DPY-6 ensures that new cuticle furrows align precisely with the previous pattern. It is possible that DPY-6 CCD does not inherently discriminate between old and new furrow collagens, but rather, it may not come into direct contact with old collagen. Given that furrow collagens are among the first to be secreted and the cuticle is layered, furrow collagens of the old cuticle may lie in a more external cortical layer during DPY-6 secretion, limiting direct interaction. So DPY-6 would position itself at the old furrow by binding to another, more basal furrow-associated component. This is further supported by the observation that, in the absence of furrow collagens, DPY-6 is still directed to a cuticular structure but is rapidly destabilized and unable to form new parallel periodic furrows.

Recent work has revealed that the apical ECM of adult *C. elegans* exhibits a collagen-based nanoscale architecture, featuring pillar-like struts within the fluid-filled layer (Adams et al., 2023). Furrow collagens are essential for the proper localization of these adult collagen struts. Intriguingly, phase separation has been proposed as a mechanism for organizing collagen struts, akin to processes observed in other ECM components, such as tropoelastin (Muiznieks et al., 2018), and predicted for *C. elegans* pharynx cuticle proteins (Kamal et al., 2022). Notably, in the absence of DPY-6, furrow collagens are still secreted into the cuticle but form large, misoriented longitudinal bundles, suggesting an intrinsic self-assembly capacity. This implies that cuticle assembly may proceed through sequential steps of guided organization and spontaneous self-assembly, integrating molecular templating with biophysical properties.

Once its job is completed, DPY-6 is removed from the matrix and does not contribute to the mature cuticle structure. Such transient factors also have been described in other matrices. For example, mammalian tooth development involves many matrix proteins that are eventually removed during the process of biomineralization (Moradian-Oldak and George, 2021). In Drosophila, a transient chitin rod shapes tracheal tube lumens (Moussian et al., 2006; Tonning et al., 2005), while transient zona pellucida (ZP) domain proteins template lens formation and sensory nanopore development before being cleared (Ghosh and Treisman, 2024; Itakura et al., 2026). Similarly, in *C. elegans*, transient pre-cuticular proteins, including multiple ZP-domain proteins, are essential for proper cuticle architecture and template varied substructures such as lateral alae ridges and sensory pores (Forman-Rubinsky et al., 2017; Fung et al., 2023; Katz et al., 2022; Serra et al., 2024). These examples underscore the key role of transient ECM components in guiding the precise spatial organization of stable matrix structures across species and tissues.

### Furrows as signaling centers

We previously showed that another CCD domain protein, SPIA-1, is also associated with furrow collagens in the cuticle (Sonntag et al., 2025). However, unlike DPY-6, SPIA-1 is not required for furrow patterning, but instead triggers immune activation upon furrow loss (Sonntag et al., 2025). Similarly, the pre-cuticular component GRL-7, a hedgehog-related protein, also localizes to furrows but appears to function in signaling rather than structural organization (Chiyoda et al., 2021; Kume et al., 2019; Serra et al., 2024). This underscores the role of furrows as sentinels, linking extracellular matrix integrity to downstream signaling pathways in the epidermis.

### How are furrows established in the first cuticle?

Interestingly, DPY-6 is dispensable for the formation of the first embryonic cuticle, raising the question of how the initial periodic pattern is established. Previous studies suggested that the actin cytoskeleton is necessary for patterning the first cuticle (Costa et al., 1997; Wang et al., 2020). During elongation, the actin cytoskeleton aligns circumferentially to constrict the epidermis, and together with muscle contraction, the embryo elongates fourfold before hatching (Lardennois et al., 2019; Vuong-Brender et al., 2017a). Pre-cuticular components are known to be crucial for elongation from the 2 fold-stage to anchor muscles and lead to efficient contraction (Kelley et al., 2015; Mancuso et al., 2012; Priess and Hirsh, 1986; Vuong-Brender et al., 2017b). Disrupting actin during elongation would have multiple effects, as elongation would cease.

We previously showed that removing the transient alignment of periodic actin does not impact furrow formation in the L4, but removing the furrow collagens abrogates actin alignment in the L4 (Aggad et al., 2023). This raises a key question: Does the epidermis pattern the matrix, or does the matrix pattern the epidermis? Further studies are required to reconcile this apparent opposition between the inside-out patterning in the embryo and the outside-in patterning in larvae. Understanding this dynamic interplay will provide deeper insights into how periodic structures are established and maintained across developmental stages.

## Materials and Methods

### Nematode strains

All *C. elegans* strains were maintained on nematode growth medium (NGM) and fed with *E. coli* OP50, as described (Stiernagle, 2006). Double mutant or KI strains were obtained by conventional crosses. Table S1A shows a list of the strains used in this study, including those previously published: CB14 *dpy-6(e14) X* (Brenner, 1974), CB5542 *dpy-6(e2762) X* (Davis et al., 2022), *BE93 dpy-2(e8) II* (Cox et al., 1980), *TLG351 dpy-6(wk113) X*, TLG350 *dpy-6(wk127) X* & TLG324 *dpy-6(wk132) X* (Lakdawala et al., 2019), XW18042 DPY-7::sfGFP (Miao et al., 2020), XW5399 *ced-1p*::NUC-1::mCherry (Li et al., 2016), *PHX3742 dpy-6(syb3742[DPY-6::mNG]) X* (Sonntag et al., 2025), PHX4550 *dpy-2(syb4550[DPY-2::mNG]) II*, PHX4583 *dpy-3(syb4583[DPY-3::mNG]) X*, PHX4549 *dpy-9(syb4549[DPY-9::mNG]) IV*, PHX4556 *dpy-10(syb4556[DPY-10::mNG]) II* and *JDW909 dpy-10(syb4556 wrd383[dpy-10::mScarlet]) II* (Ragle et al., 2026).

### KI lines

All the knock-in, deletion or mutant strains were designed relying on information from WORMBASE (Davis et al., 2022) and were made by SEGiCel (SFR Santé Lyon Est CNRS UAR 3453, Lyon, France) by CRISPR/Cas9 and were confirmed by PCR genotyping using primers outside the homology arms and Sanger sequencing, sequences are available upon request. MCP1001 *dpy-6(syb3742[DPY-6ΔCCD::mNG]) X* is a deletion of the CCD from L1200 to the end of DPY-6 in the strain PHX3742 *dpy-6(syb3742(DPY-6::mNG)) X* (Sonntag et al., 2025).

MCP836 *dpy-8(bab836[mNG_DPY-8]) X* is an mNG insertion N-ter of DPY-8.

MCP858 *dpy-8(bab858[DPY-8::mNG]) X* is an mNG insertion at position S394 in DPY-8.

MCP992 *dpy-8(bab858[DPY-8::mNG]) X* is a mutation of the furin site in AxxA in the MCP858 strain.

MCP914 *dpy-7(bab914[mNG::DPY-7]) X* is an mNG insertion N-ter of DPY-7.

MCP861 *dpy-7(bab861[DPY-7::mNG]) X* is an mNG insertion at position G76 in DPY-7.

MCP990 *dpy-7(bab861[DPY-7::mNG]) X* is a mutation of the furin site in AxxA in the MCP861 strain.

### RNA interference

RNAi bacterial clones were obtained from the Ahringer or the Vidal libraries (Kamath et al., 2003; Rual et al., 2004) and verified by sequencing (see Table S1B). RNAi bacteria were seeded on NGM plates supplemented with 100 g/ml ampicillin and 1 mM Isopropyl-β-D-thiogalactopyranoside (IPTG). Worms were transferred onto RNAi plates as L1 larvae and cultured at 25 °C until L4 or young adult stage. For bli-4 RNAi experiment, L3 larvae were transferred from OP50 plate onto RNAi plate and imaged at L4 stage. For pulse experiment, L4 larvae were put in *bli-4* RNAi for 3 hours, then transferred back on OP50 and then transferred onto a new OP50 plate the next morning; the next generation from this second plate was analyzed (Fig S2B-C). In all our experiments, we used *sta-1* as a control, as we have shown over the last decade that it does not affect the development nor any stress or innate response in the epidermis (Dierking et al., 2011; Lee et al., 2018; Taffoni et al., 2020; Zhang et al., 2021; Zugasti et al., 2014).

### Atomic force microscopy

As previously described (Aggad et al., 2023), the worms were paralyzed in 15 mg/mL 2,3-butanedione monoxime (BDM, Sigma) for 10-30 min. They were then aligned on a ∼2-mm-thick 4 % freshly prepared agarose pad in a Petri dish (30 mm diameter). For full immobilization, the worms were fixed (head and tail) with 2 % low-melting agarose (GibcoBRL) instead of glue, as we found that the latter altered the cleanliness of the topographic image. After the agarose dried, we immersed again the worms in the BDM solution. AFM data were obtained using a NanoWizard4 (Bruker-JPK) during up to one hour following mounting. 10×10 µm areas on top of the worms were scanned using PFQNM-LC-cal cantilevers (Bruker) in quantitative imaging mode. The inverse of the optical lever sensitivity (invOLS) was calibrated from the thermal spectra in liquid, using the pre-calibrated spring constant of the cantilevers and the correction factors described in (Rodriguez-Ramos and Rico, 2021). The force setpoint was set at 0.5 nN, speed at 50 µm/s and the range at 100 nm in Z. All topographical images (256×256 px) are flattened using the plane fitting and line levelling options of the JPK DP software to correct for sample tilt.

### Confocal microscopy

Worms were staged into distinct substages during the 12-hour L4 stage using vulval morphology as a proxy for developmental timing, as described previously (Aggad et al., 2023; Cohen et al., 2020; Mok et al., 2015). They were mounted on a 2 % agarose pad, in a drop of 1 mM levamisole in 50 mM NaCl. Images were acquired during the following 60 min, using Zeiss confocal laser scanning microscopes (LSM780, 880 or 980) and the acquisition software Zen with a Plan-Apochromat Oil DIC M27 40×/1.4 objective. Pinhole size was set to 1 AU. Samples were illuminated with 488 nm (sfGFP, mNG) and 561 nm (mScarlet, mCherry) with varied laser power based on protein abundance and tissue imaged, with 4 lines accumulation and 800 gain settings. Spectral imaging combined with linear unmixing was used to separate the autofluorescence of the cuticle. Images in Fig. 3F-H were collected as described in (Birnbaum et al., 2023). In all images, only one confocal plane is presented, unless stated differently. Images were analyzed and processed in Fiji (Schindelin et al., 2012). To quantify fusion protein accumulation in the extraembryonic space, fluorescence intensity was measured in a 3 × 3 μm region within a single medial confocal slice of each specimen, as described in (Birnbaum et al., 2023).

### Imaris 3D reconstitution

Three-dimensional reconstruction of the signal was performed using confocal image stacks acquired with Zen software. Z-stacks were captured at 0.39 µm intervals with 2-line accumulation during confocal microscopy. The signal of interest was segmented using the Imaris Machine Learning module: a supervised classifier was trained on manually annotated regions to distinguish signal from background and applied to the full volume to generate a voxel-wise probability map. A 3D surface was then reconstructed using the Imaris Surface Creation tool, with automatic thresholding based on the model-generated probabilities. Smoothing and minimum object size parameters were optimized to eliminate artifacts.

### Dumpy phenotype validation

L4 larvae were selected one day prior to analysis and cultured at 20°C until they reached Day 1 adulthood. Worms were then imaged using AxioVision software with a calibrated scale. Worm lengths were measured in Fiji (Schindelin et al., 2012) using the “Segmented Line” tool, tracing a line from head to tail along the curvature of the worm.

### Statistical analysis

Data were analyzed using GraphPad Prism 10.3. In all dot plots, lines and error bars represent the median and range, respectively, with each dot corresponding to a measurement from a single animal. Normality was assessed using the Shapiro-Wilk test, followed by ordinary one-way ANOVA with Tukey’s post hoc test for multiple comparisons. Differences were considered statistically significant at p < 0.05.

## Supporting information

Supplemental informations

## Acknowledgements

We thank Jordan Ward & Andrew Chisholm for providing the DPY-2::mNG, DPY-3::mNG, DPY-9::mNG, DPY-10::mNG and DPY-10::mScarlet strains prior to publication, Jonathan Hodgkin and Tina Gumienny for *dpy-6* alleles, Mathieu Fallet for his assistance with image quantifications. We thank Margaux Gibert and SEGiCel (SFR Santé Lyon Est CNRS UAR 3453, Lyon, France) for the elaboration of all the different tagged and mutated version of DPY-7::mNG and DPY-8::mNG strains, as well as the mutated DPY-6ΔCCDmNG. Some *C. elegans* strains were provided by the CGC, funded by NIH Office of Research Infrastructure Programs (P40 OD010440). We thank the staff at WormBase for their outstanding community work, including the maintenance of a curated database (Davis et al., 2022), Michel Labouesse and Jonathan Ewbank for discussions and comments on the manuscript. We acknowledge the PICSL imaging facility of the CIML (ImagImm), a member of the national infrastructure France-BioImaging supported by the French National Research Agency (ANR-24-INBS-0005 FBI - BIOGEN), Région SUD, IBiSA and Amidex fundation.

## Supplementary Materials

**Fig S1. Embryonic and larval expression of mNG tagged furrow collagens and DPY-6**.

(A) Representative confocal images of mNG-tagged furrow collagen and the DPY-6 proteins (green signal), merged with the transmission channel, during embryonic development and L1 larvae stage; n>5 for each stage; scale bar, 5 µm. Note that for DPY-10::mNG in early L1 and DPY-6::mNG in late L1, the worm is not lying on its side, unlike the other L1 images where the demarcation of the seam cells is visible.

(B) Representative confocal images of DPY-7::mNG (green) and FBN-1::mCherry (magenta) in L4.7 worm and the overlay of both signals ; n>10; scale bar, 5 µm.

**Fig S2. Phenotypes of different *bli-4* inactivation**.

(A) Different time and duration of *bli-4* RNAi result in diverse phenotypes, ranging from 100% larval lethality to viable Dpy phenotype. Yellow and orange box represent time on OP50 plate or on *bli-4* RNAi plate, respectively.

(B) Pie chart representing the observed phenotype of progeny from DPY-8::mNG L4 larvae subjected to a 3 hour pulse of *bli-4* RNAi (as F1), and progeny of the severe dumpy F1 worms (as F2).

(C) Dpy L4 larvae progeny from DPY-8::mNG L4 larvae subjected to a 3 hours pulse of *bli-4* RNAi (same worm as Fig.3E); red drawings represent the bifurcated alae running above the seam cell along the worm; scale bar, 30 µm for the main image, 5 µm for the zoom images.

(D) Representative confocal images of DPY-7::mNG in L3 larvae grown on RNAi plates. Inactivation of *kpc-1* alone does not affect furrow collagen secretion or positioning. Combined *kpc-1* and *bli-4* RNAi also fails to exacerbate the *bli-4* phenotype; n > 10; scale bar, 10 µm.

**Supplementary table 1**

This table includes all the worm strains and RNAi clones used in this study.

**Supplementary table 2**

This table includes all the raw data for all the quantifications.

