## Supplemental informations for "Pre-cuticle DPY-6 acts as a blueprint for aECM periodic organization in *C. elegans*"

### Figure S1

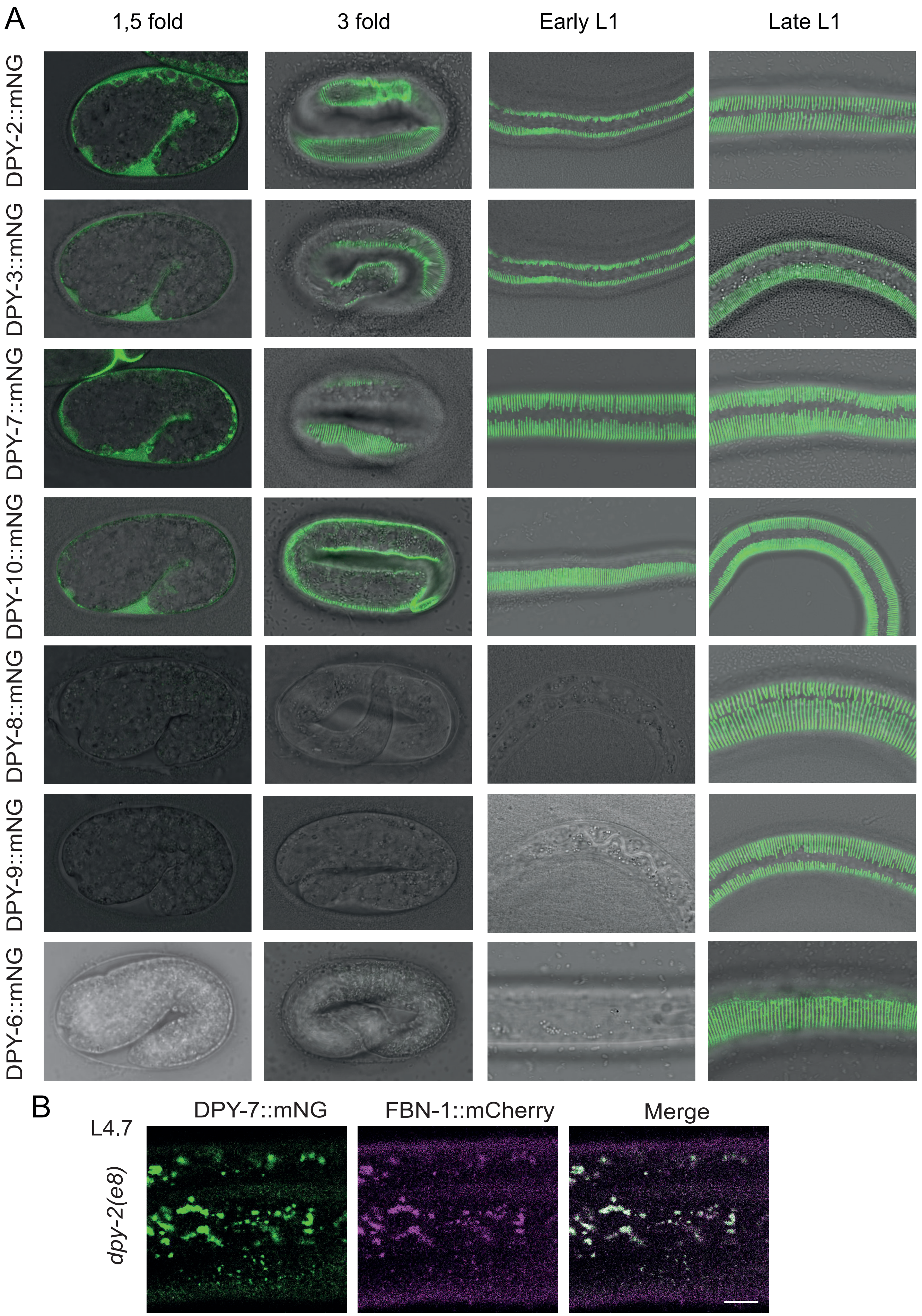

A
1,5 fold
3 fold
Early L1
Late L1
DPY-2::mNG
DPY-3::mNG
DPY-7::mNG
DPY-10::mNG
DPY-8::mNG
DPY-9::mNG
DPY-6::mNG
B
DPY-7::mNG
FBN-1::mCherry
Merge
L4.7
*dpy-2(e8)*

Figure S2

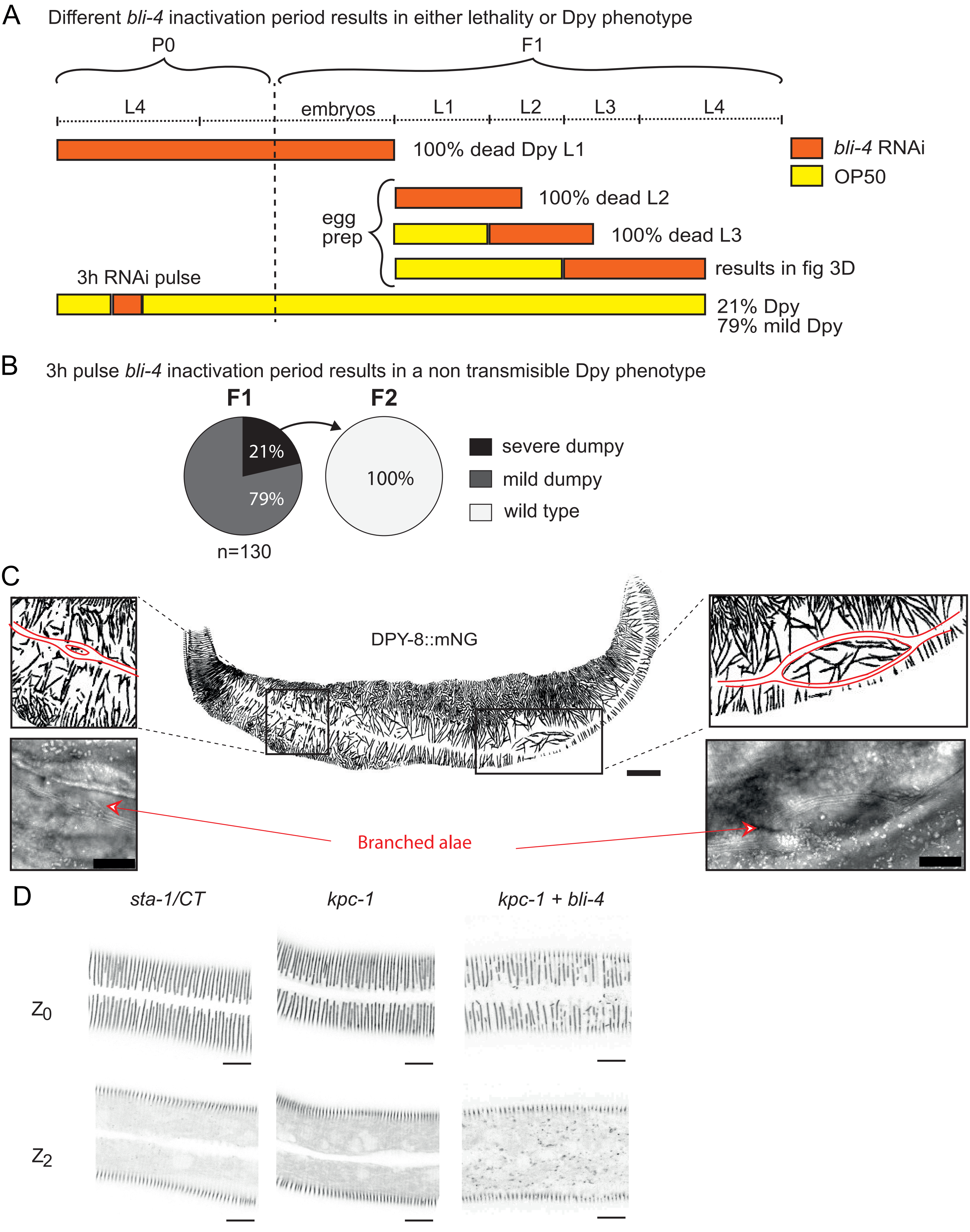
A
Different bli-4 inactivation period results in either lethality or Dpy phenotype
P0
F1
L4
embryos
L1
L2
L3
L4
100% dead Dpy L1
bli-4 RNAi
OP50
egg prep
100% dead L2
100% dead L3
results in fig 3D
3h RNAi pulse
21% Dpy
79% mild Dpy
B
3h pulse bli-4 inactivation period results in a non transmisible Dpy phenotype
F1
F2
21%
79%
100%
n=130
severe dumpy
mild dumpy
wild type
C
DPY-8::mNG
Branched alae
D
sta-1/CT
kpc-1
kpc-1 + bli-4
Z0
Z2

### Table S1a: *C. elegans* strains

| Strain | Genotype | Reference | Figure |
|---|---|---|---|
| UP4108 | *bli-4(cs281); qxIs722[dpy-7p::dpy-7::SfGFP] II* | Birnbaum et al, 2023 | 3G,H |
| UP4111 | *bli-4(cs302) I; qxIs722[dpy-7p::dpy-7::SfGFP] II* | Birnbaum et al, 2023 | 3F |
| JDW909 | *dpy-10(syb4556 wrd383[dpy-10::mScarlet]) II* | Ragle et al, 2025 | 1C, 5B,E |
| PHX4556 | *dpy-10(syb4556[DPY-10::mNG]) II* | Ragle et al, 2025 | 1B,C,D, 1SA |
| IG2250 | *dpy-2(e8) II; dpy-6(syb3742[DPY-6::mNG]) X* | this study | 5G |
| PHX4550 | *dpy-2(syb4550[DPY-2::mNG]) II* | Ragle et al, 2025 | 1B,C,D, 1SA |
| PHX4583 | *dpy-3(syb4583[DPY-3::mNG]) X* | Ragle et al, 2025 | 1B,C,D, 1SA |
| CB14 | *dpy-6(e14) X* | Brenner et al, 1974 | 4G-H, 5A |
| CB5542 | *dpy-6(e2762) X* | Simmers et al, 2003 | 4F,H |
| PHX3742 | *dpy-6(syb3742[DPY-6::mNG]) X* | Sonntag et al, 2025 | 4C, 1SA |
| IG2232 | *dpy-6(syb3742[DPY-6::mNG]) X; aaaIs12[fbn-1p::FBN-1::mCherry; ttx-3p::GFP] V* | this study | 4E |
| IG2244 | *dpy-6(syb3742[DPY-6::mNG]) X; dpy-10(syb4556 wrd383[DPY-10::mScarlet]) II* | this study | 4B |
| IG2258 | *dpy-6(syb3742[DPY-6::mNG]) X; qxIs257[ced-1p::NUC-1::mCHERRY, unc-76(+)] V* | this study | 4D |
| MCP1001 | *dpy-6(syb3742[DPY-6ΔCCD::mNG]) X* | this study | 4F,G,H |
| IG2247 | *dpy-6(syb3742[DPY-6ΔCCD::mNG]) X; dpy-10(syb4556 wrd383[DPY-10::mScarlet]) II* | this study | 5D,F |
| TLG351 | *dpy-6(wk113) X* | Lakdawala et al, 2019 | 4F,H |
| TLG350 | *dpy-6(wk127) X* | Lakdawala et al, 2019 | 4F,H |
| IG2245 | *dpy-6(wk127) X; dpy-10(syb4556 wrd383[DPY-10::mScarlet]) II* | this study | 5B,E |
| TLG324 | *dpy-6(wk132) X* | Lakdawala et al, 2019 | 4F,G,H |
| IG2246 | *dpy-6(wk132) X; dpy-10(syb4556 wrd383[DPY-10::mScarlet]) II* | this study | 5B |
| MCP990 | *dpy-7(bab861[DPY-7(AXXA)::mNG]) X* | this study | 2A,B,C,F,G |
| IG2256 | *dpy-7(bab861[DPY-7(AxxA)::mNG]) X ; qxIs257[ced-1p::NUC-1::mCHERRY, unc-76(+)] V* | this study | 2E |
| MCP861 | *dpy-7(bab861[DPY-7::mNG]) X* | this study | 1B,C,D, 2A,B,C,D,F |
| IG2252 | *dpy-7(bab861[DPY-7::mNG]) X; dpy-2(e8) II; aaaIs12[fbn-1p::FBN-1::mCherry; ttx-3p::GFP] V* | this study | S1B |
| IG2255 | *dpy-7(bab861[DPY-7::mNG]) X; dpy-2(e8) II; qxIs257[ced-1p::NUC-1::mCHERRY, unc-76(+)] V* | this study | 1E |
| MCP914 | *dpy-7(bab914[mNG::DPY-7]) X* | this study | 2A,B,C,D |
| IG2254 | *dpy-7(bab914[mNG::DPY-7]) X; qxIs257[ced-1p::NUC-1::mCHERRY, unc-76(+)] V* | this study | 2E |
| MCP836 | *dpy-8(bab836[mNG_DPY-8]) X* | this study | 2A,B,C |
| MCP992 | *dpy-8(bab858[DPY-8(AXXA)::mNG]) X* | this study | 2A,B,C |
| MCP858 | *dpy-8(bab858[DPY-8::mNG]) X* | this study | 1B,D, 2A,B, 3B,C,D,E |
| PHX4549 | *dpy-9(syb4549[DPY-9::mNG]) IV* | Ragle et al, 2025 | 1B,D, 1SA |
| XW5399 | *qxIs257[ced-1p::NUC-1::mCHERRY, unc-76(+)] V* | Li et al, 2016 | cross |
| XW18042 | *qxSi722[dpy-7p::DPY-7::sfGFP; ttTi5605] II* | Miao et al, 2020 | 3F,G,H |
| N2 | Wild-type | Brenner, 1974 | 2B, 4G,H |

### Table S1b: RNAi clones

| **Gene** | **Clone** |
|---|---|
| *sta-1* | sjj_Y51H4A.o |
| *bli-4* | mv_K04F10.4 |
| *kpc-1* | sjj_F11A6.1 |
| *dpy-6* | sjj_F16F9.2 |
| *dpy-2* | sjj_T14B4.6 |
| *dpy-3* | sjj_EGAP7.1 |
| *dpy-7* | sjj_F46C8.6 |
| *dpy-8* | sjj_C31H2.2 |
| *dpy-9* | sjj_T21D12.2 |
| *dpy-10* | sjj_T14B4.7 |
